# A model of collective behavior based purely on vision

**DOI:** 10.1101/589663

**Authors:** Renaud Bastien, Pawel Romanczuk

## Abstract

Classical models of collective behavior often take a “birds-eye perspective,” assuming that individuals have access to social information that is not directly available (e.g., the behavior of individuals outside of their field of view). Despite the explanatory success of those models, it is now thought that a better understanding needs to incorporate of the perception of the individual, i.e. how internal and external information are acquired and processed. In particular, vision has appeared to be a central feature to gather external information and influence the collective organization of the group. Here we show that a vision based model of collective behavior is sufficient to generate organized collective behavior in the absence of spatial representation and collision. Our work suggests a novel approach for development of purely vision-based autonomous swarm robotic systems, and formulates a mathematical framework for exploration of perception-based interactions and how they differ from physical ones. Thus, it is of broader relevance for self-organization in complex systems, neuroscience, behavioral sciences and engineering.

---

Models of collective behaviour often rely on phenomenological interactions of individuals with neighbors^1–4^. However, and contrary to physical interaction, these social interactions do not have a direct physical reality, such as gravity or electromagnetism. The behavior of individuals is influenced by their representation of the environment, acquired through sensory information. Current models often suggest that individuals are responding to the state of movement of their neighbors – their (relative) positions and velocities – which are not explicitly encoded in the sensory stream. Thus such phenomenological interactions implicitly assume internal processing of the sensory input in order to extract the relevant state variables. On the other hand, neuroscience has made tremendous progress in understanding various aspects of the relation of sensory signals and movement response, yet connections to large-scale collective behavior are lacking. Although evidence has been found for neural representation of social cues in the case of mice^5^ and bats ^6^, yet details and role of these internal representations remain unclear, in particular in the context of coordination of movement. Collective behavior crucially depends on the sensory information available to individuals, thus ignoring perception by relying on ad-hoc rules, strongly limits our understanding of the underlying complexity of the problem. Besides, it obstructs the interdisciplinary exchange between biology, neuroscience, engineering, and physics.

Recently, the visual projection field has appeared as a central feature of collective movements, in fish^7–10^, birds^11^ or humans^12^. Due to the geometrical nature of vision, *i.e*. the projection of the environment, vision appears as a good starting point to explore the relationship between sensory information and emergent collective behaviors. Some models have attempted to relate vision and movement ^8, 11, 13, 14^. However, they use vision as a motivation to refine established social interaction models or rely on additional interactions based on information not explicitly represented in visual input such as distance or heading direction of the neighboring individual.

Here, we propose a radically different approach by introducing a general mathematical framework for purely vision-based collective behavior. We use a bottom-up approach employing fundamental symmetries of the problem to explore what types of collective behavior may be obtained with as minimal as possible requirements.

Formally, we can write the movement response of an agent to the visual field in three spatial dimensions as the following evolution equation for its velocity vector **v**_**i**_:

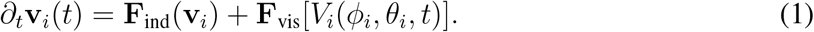

The first term accounts for the self-propelled movement of an individual. Here, we use a simple linear propulsion function: 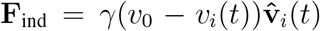, with *v*_0_ being the preferred speed of an individual, *γ* the speed relaxation rate, and 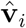 the heading direction vector of the focal individual with 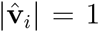. The second term accounts for the movement response to the instantaneous visual sensory input given by the visual field *V*_*i*_(*ϕ*_*i*_, *θ*_*i*_, *t*), experienced by the individual *i*. *ϕ*_*i*_ and *θ*_*i*_ are the spherical component relative to the individual *i* and **F**_vis_ is an arbitrary transformation of the visual field. This function does not have an explicit dependence on the other individual properties.

The physical, visual input corresponds to a spatio-temporal intensity and frequency distribution of the incoming light. In our framework, we consider *V* to be an abstract, arbitrary representation of the visual input. In particular, *V* can implicitly account for relevant sensory (pre-)processing, e.g. it can represent colors or brightness of the visual scene. Furthermore, *V* can also account for higher order processing of visual stimuli such as object identification and classification. Equation 1 describes the projection of the full information encoded in the visual field onto the low dimensional movement response and must hold for any particular choice of visual field.

**Figure 1:**
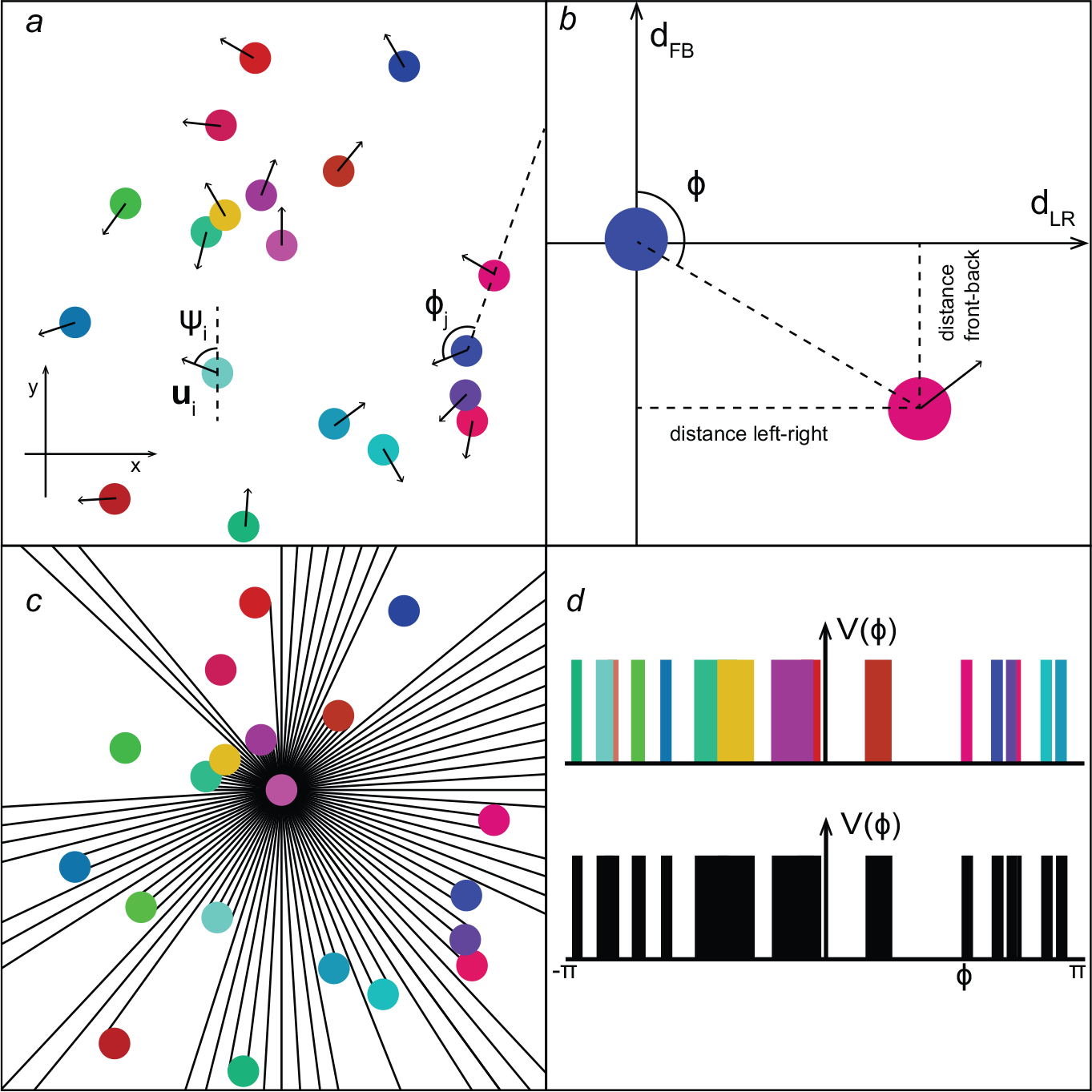
a. A set of disks with diameter *D* is considered. Each disk is propelled in the direction *ψ*_*i*_ with a velocity *v*_*i*_(*t*). b. A co-moving referential can be defined following the movement of the disk *j*. This referential is centered on the position of the disk, (*x*_*j*_, *y*_*j*_), and oriented so that the vertical axis is aligned with the direction *ψ*_*j*_. The position of other objects can be recovered through their left-right, *d*_*LR*_ and front-back distances *d*_*FB*_ relative to the the disk *j*. *ϕ* represents the swiping angle. c. Representation of the visible field of the pink disk, through ray casting. The position of the eye is considered to be at the center of the disk with fully circular point-of-view, *i.e*. no blind angles. d. The projection of the visual field in 2D is given by a 1D function. On top, objects can be represented by their colors. However on the bottom part, a binary visual field is given. It is not possible to distinguish individuals.

In order to simplify the description, we limit our analysis here to the two-dimensional case. Without any loss of generality, **F**_vis_ can be written as

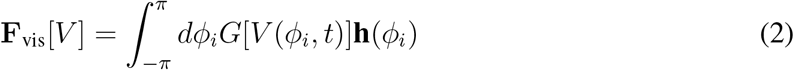

The functional *G*[*V*], encodes what information from the visual field influences the movement response and how. An arbitrary *G* can be expanded as a series of derivative in space and time and power series of the visual field. This account for any function of the visual field; e.g. specific functions of the visual cortex such as detection of edges in all directions or optical flow. The function **h**(*ϕ*_*i*_): ℝ *→* ℝ^2^, on the other hand, encodes the fundamental properties of the perception-motor system (“the observer”) independent on the specific visual input, e.g. symmetries of the movement response, or spatial dependence of perception (e.g. blind angle). Experimental data in fish have shown that the variation of orientation depends on the left-right position of the other individual, while variations of speed depend on front back position. The components of **h** are therefore expanded as a Fourier series in *ϕ*.

Up to this point no approximation has been made, the model is then as general as possible regarding response to an arbitrary visual field. In order to develop a systematic understanding of how collective behavior can arise from the visual field, we propose first a minimal model of vision based interactions. First, we assume that individuals respond to an instantaneous, binary visual field, i.e. the visual projection field *V* (*φ, t*) only accounts for the presence or absence of objects and no other properties. Second, we consider an expansion of an arbitrary functional *G* in terms of the lowest order of space and time derivatives in *V*. The velocity vector of an agent in 2D is determined by the velocity with respect to the heading direction *v*_*i*_(*t*) and the polar angle determining the heading vector *ψ_i_*(*t*). The simplest equations of movements, satisfying the fundamental symmetries from ^15^, read:

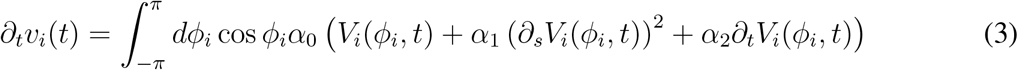

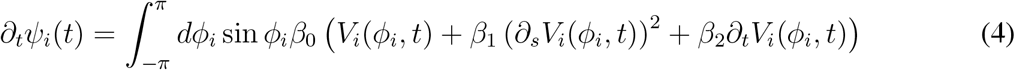

The first terms in the brackets describe the movement response to the perceived “mass” of the objects in the visual projection, the second describe the response to edges, while the third ones account for dynamical changes such as translation or loom. The coefficients *α*_*m*_ and *β*_*n*_ are arbitrary constants obtained from the expansion of *G*. In the following, we show that coordinated collective movement can also emerge without considering temporal derivatives, i.e. by setting *α*_2_ = *β*_2_ = 0.

The first terms associated with the mass of the visual field creates a short-range interaction that decreases as the object gets further. On the contrary, the second terms with the first derivative with respect to the visual field coordinate, yield long-range interaction due to the non-linearity of the sin/cos function. Thus, these lowest order terms, neglecting temporal derivative, are sufficient to generate short-range repulsion and a long-ranged attraction: The individual is repelled by the mass of the object on its visual field while getting attracted by the edges. Based, on the choice of corresponding interaction parameters, we can define an equilibrium distance, where attraction and repulsion balances (see Supp. Information for details). This equilibrium introduces now a characteristic metric length scale into the system, despite the lack of any representation of space at the level of individual agents.

**Figure 2:**
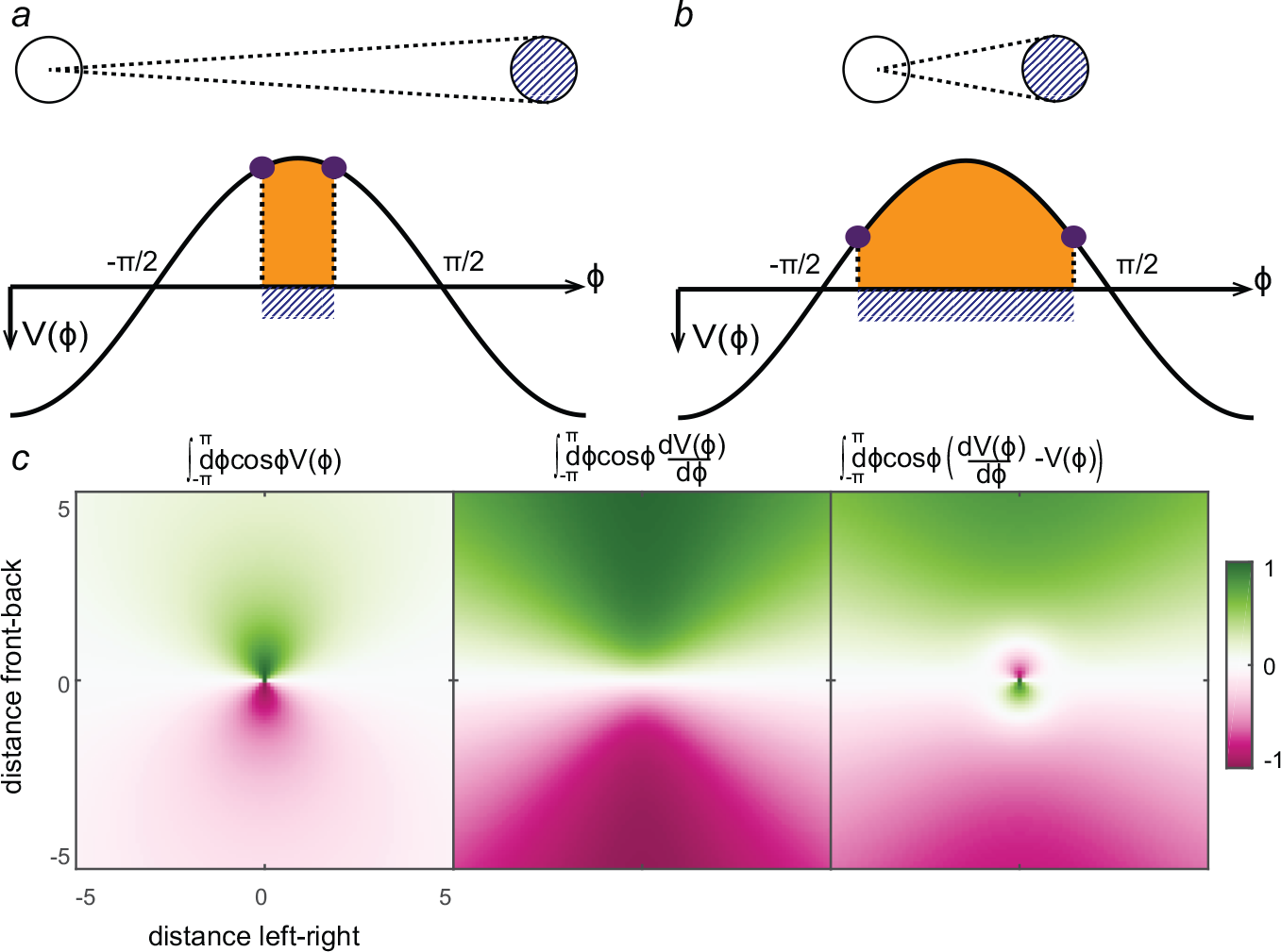
Effects of the terms of the equations 3 on a focal observer according to the relative position an other disk. The white disk is looking straight at the blue disk with an eye positioned in the center. When the object is far, a, the mass of object on the projection of the visual field, *V* (*ϕ*) is smaller than when the object is close, b. When integrating with a cosine function, the mass of the object (in orange) results in a larger integration for closer object, while the edges (in purple) sum larger element of the cosine. c. For different relative positions between both disks, the mass of the object produces a short range interaction, while the edges creates a long range interaction. The difference of those two terms can create a short range repulsion (with the mass of the object)/ long range attraction (with the edges of the object).

A systematic exploration of the collective behavior of multiple agents interacting through the minimal vision model reveals the emergence of cohesive, collective behaviors for a wide range of parameters and different group sizes (Figs 3, 4). In particular, we observe robust self-organized collective movements, emerging from the interplay of visual perception and the movement response of individuals. The degree of coordination can be quantified through the normalized average velocity of the group also referred to as orientational order or polarization (Fig. 4). Besides the ability to exhibit ordered, directed collective movement, an often neglected property of collective movement, is the ability of agents to avoid collisions. This might be in particular critically important for artificial swarm robotics systems. Here, we can identify extended regions of parameter space without any collisions overlapping with the regions of ordered motion.

**Figure 3:**
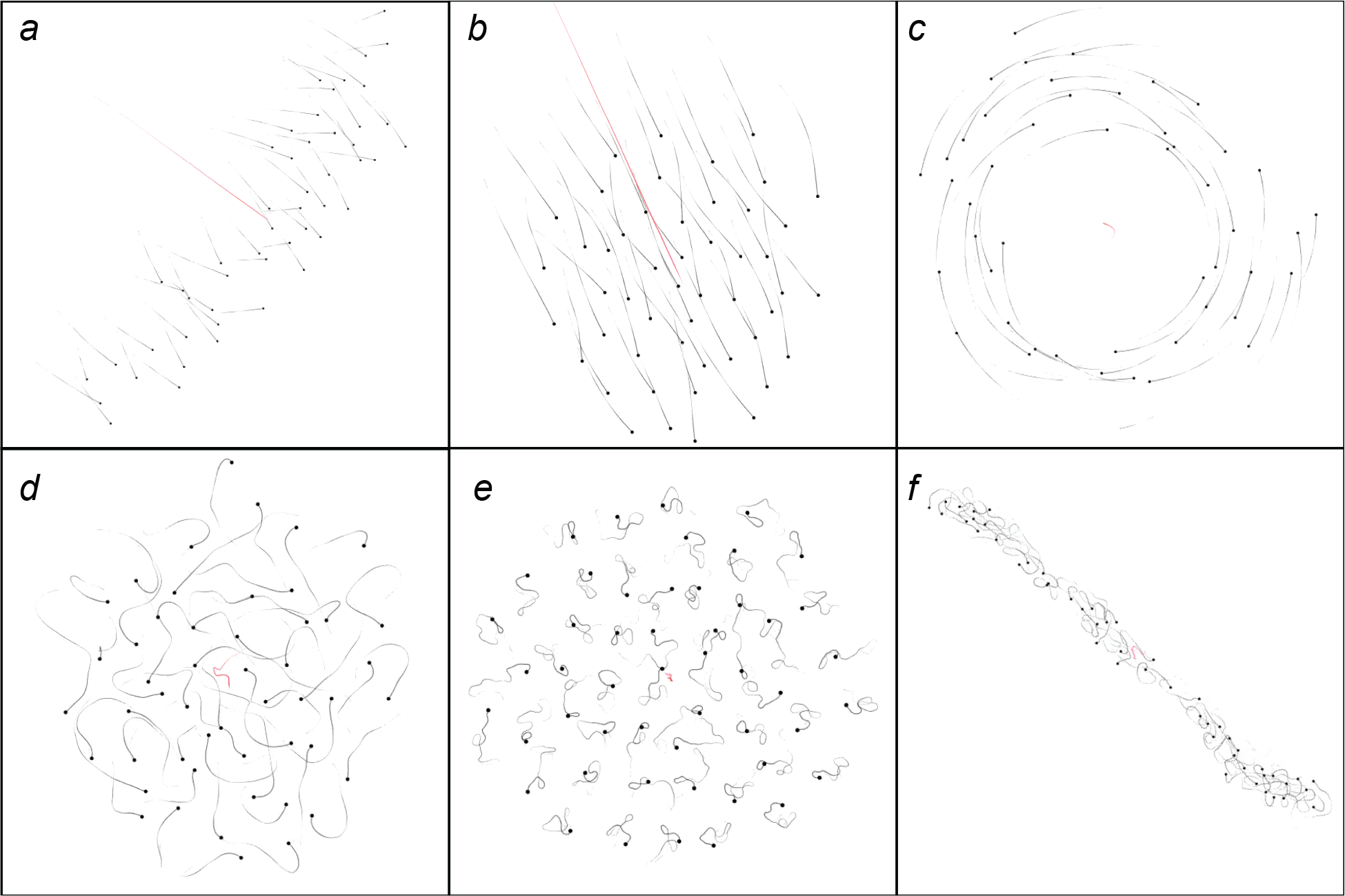
Different collective behaviours observed in the model for N=50 individuals. a. Polarized on a line perpendicular to the movement. b. Polarized in a circular shape. c. rotating, no preferred direction is chosen here, so individuals are turning in both directions at the same time. d. Swarm behaviour where individuals are moving freely in the swarm. e. Crystal-like configuration. f. Tube-like configuration.

**Figure 4:**
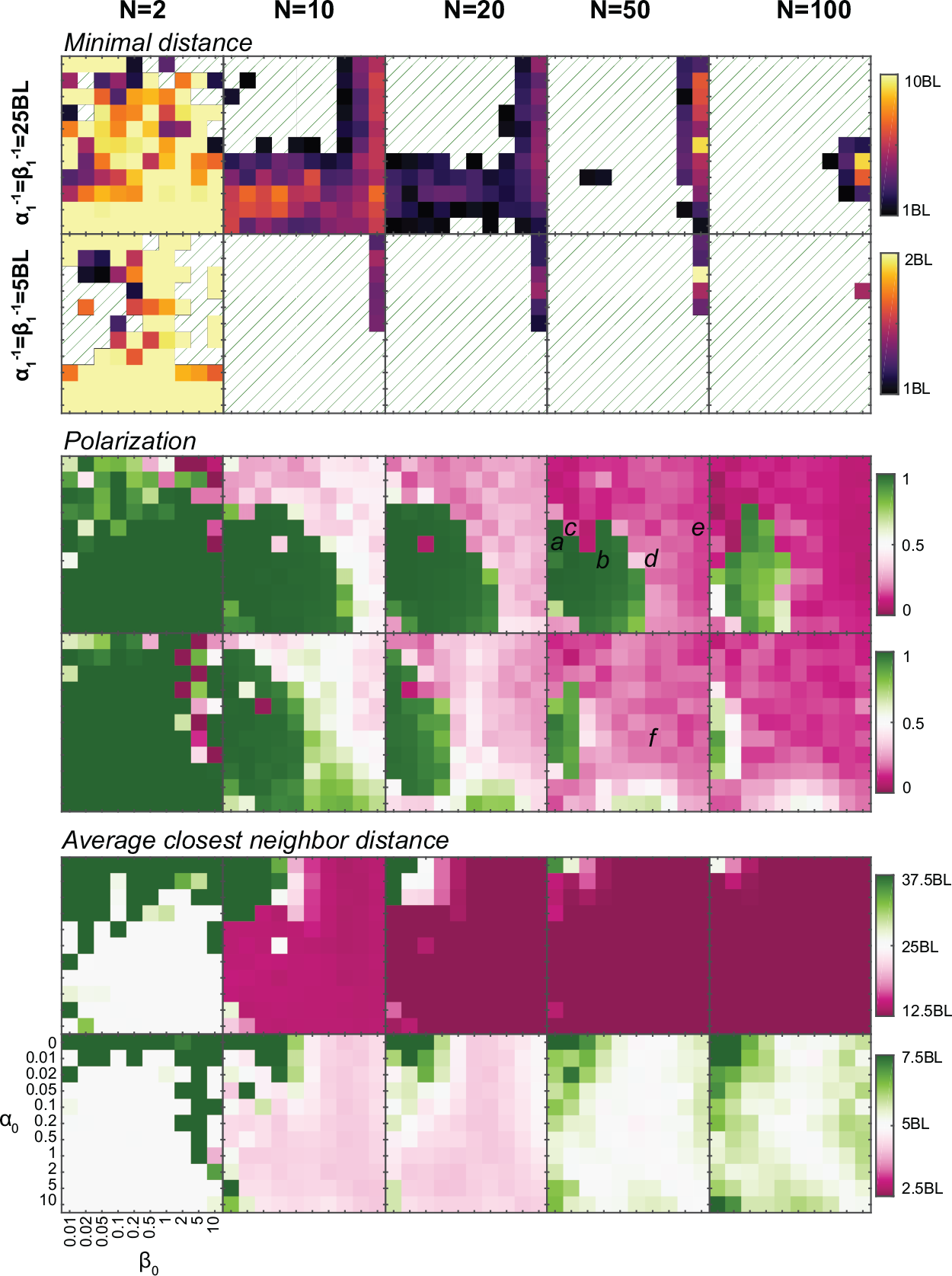
From top to bottom, the minimal distance observed in the simulations^17^, the polarization of the swarm and the average closest neighbor distance as function of *α*_0_ and *β*_0_, for different numbers of individuals, from left to right *N* = 2, 10, 20, 50 and 100, and for two different values of the equilibrium distance, top row *α*_1_ = *β*_1_ = 25*BL* and bottom row *α*_1_ = *β*_1_ = 5*BL*. For the minimal distance, dashed lines represents distance that are less than one body-length, so the objects are colliding.

The observation of coordinated motion without any collisions is, in particular, remarkable as our minimal vision model does not take any time-derivatives of the visual field (i.e. optical flow) into account, and thus lacks any explicit or implicit alignment mechanisms (see e.g. ^2, 16^). Furthermore, individuals do not know where they are relative to others; thus they do not use any information on the number or the distance of other individuals.

By relating perception and movement response, we have shown that a simple model of collective behavior can be constructed directly without the need to explicit ad-hoc rules of coordination between individuals. This model does not specify spatial representation, explicit alignment or explicit representation of other individuals. Those features are then not necessary features of organized collective behavior. It is important to emphasize this last point. If this model could be seen as a simple rewriting of classical models in the referential of an individual, it also calls into question the underlying representations that are used in classical models. Can animals identify the positions in space of other individuals? How many neighbors can be represented in the vision of an individual? The answer to this question should arise from neurophysiological data, but their link to the movement response needs to be explicitly stated. Furthermore, a proper spatial scale is introduced in the system through the size of the animal and not through some parameters in the equation. That opens the way to study effects such as anisotropy which was not possible before.

We believe the model framework is also of relevance to physics and dynamical systems, as on the one hand, it is a paradigmatic example of a class of models where interaction between individuals are not based on physical force fields, but solemnly on the perception and internal representation of the social environment by the local agent. In our specific case, the coupling between agents is based on a lower dimensional projection of the actual dynamical behavior of many agents, which at the same time respond to their perceptional input. In contrast to well studied force-based models in dynamical systems and statistical physics, here we even appear to lack the mathematical tools to systematically explore the emergent collective behaviors for this novel type of models.

Furthermore, the simple vision-only interaction discussed here has some interesting properties. It does not correspond to a simple superposition of binary interactions and does not rely on arbitrary cutoffs or thresholds. Thus it results in a self-consistent description of interactions from a single individual up to large groups, naturally accounting for effects like self-organized marginal opacity ^11^ due to saturation of the visual field.

Finally, the absence of collision in such models should open ways to create artificial robotics systems, such as a swarm of drones, where the only input of the system is fully decentralized, based on perception and not through the explicit transfer of information between constituting agents.

## Methods

### 1 Model Construction

We start with equation 1. To simplify the calculations, it is assumed that the movement is going to be contained inside a single plane. The velocity of an individual *i*, **v**_**i**_, is then described by its direction *ψ*_*i*_ and its magnitude *v*_*i*_.

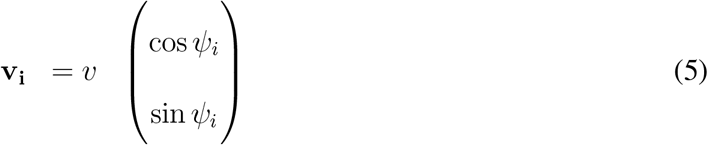

It is then possible to easily define elementary vectors relative to the orientation of the individual *i*

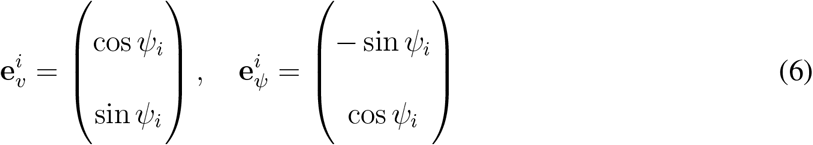

so that

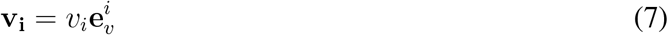

The equation 1 can be split in two equations

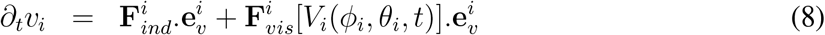

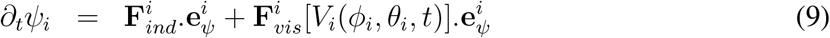

that describes the variation of magnitude *u* and the variation of direction respectively. Now that the basics have been defined, the model and its hypotheses are described. First, as the individual behavior is not the first focus of the model, and in order to prevent the model from diverging, the individual behavior **F**_*ind*_ is siply given by

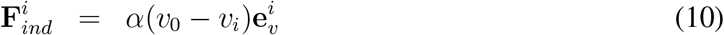

where *α* is some constant, and *v*_0_ the preferred velocity magnitude of the individuals. As the velocity vector of the individual does not depend directly on *ϕ*, 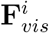 can be rewritten as an integral over the visual space

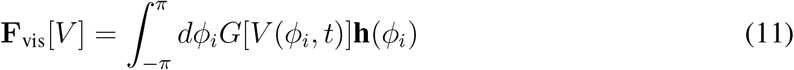

Here, **h**(*ϕ*) is an arbitrary vector function, determining the projection of *G*[*V*] on the low-dimensional movement response. It can be seen as a target function for the visual input. The symmetries of the system can be inferred from^15^. Here, they measure the values of *∂*_*t*_*v*_*i*_ and *∂*_*t*_*ϕ*_*i*_ for one individual, as a function of the relative position of another individual. *∂*_*t*_*ϕ*_*i*_ is anti-symmetric between left and right whereas *∂*_*t*_*v*_*i*_ is anti-symmetric between front and back. The target functions **h**(*ϕ*) carries those symmetries and can be rewritten as

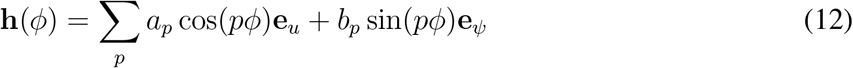

Furthermore, the assumption of **h**(*ϕ*) component along **e**_*u*_ and **e**_*ψ*_ being symmetric and antisymmetric functions around *ϕ* = 0, respectively, is required to ensure the absence of permanent rotational motion of individual agents.

Inserting Eq. 12 into Eq. 11 eventually yields, the movement equations:

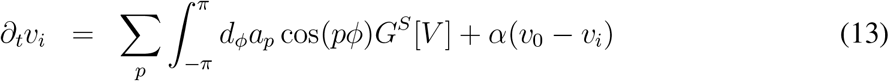

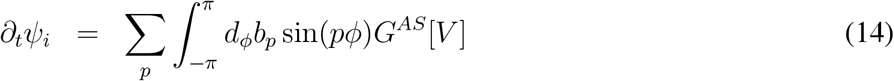

Here, we split the function *G*[*V*] into its symmetrical part, *G*^*S*^[*V*], and its anti-symmetrical part *G*^*AS*^[*V*],

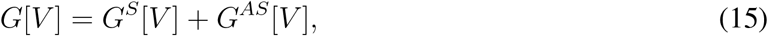

based on the non-zero contribution to the above integrals. The movement is then driven by the discrepancies in the symmetry of the visual field. The asymmetry between left and right will modify the direction of the individual while the asymmetry between front and back will modify the magnitude of the velocity.

Now we can rewrite the symmetric and anti-symmetric parts in terms of derivatives in time, 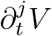, and space, 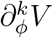:

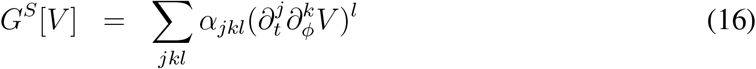

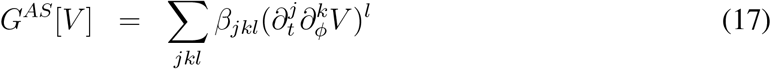

Until now we have been as general as possible. The visual field remains here an abstract function that needs to be appropriately defined. A tremendous diversity of parameters and orders can be considered. However, as we now have described the general model, it is possible to study each model independently, starting with the simplest one, and to extract the mechanisms expected at each scale. The simplest description for the visual field is then given by the following assumptions

**A1** The visual field is binary, when an object is in the field *V* = 1, and if not *V* = 0 (no complex sensitivity based on distance…)

**A2** There is no complex cognitive function (no hierarchy between individuals, no individual identification, no selective attention, etc)

If we only consider the first order of the Fourier transform, the full model for a binary visual field is given by

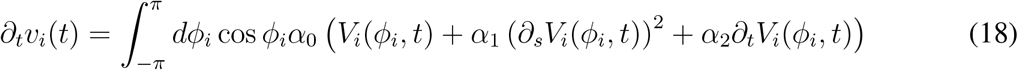

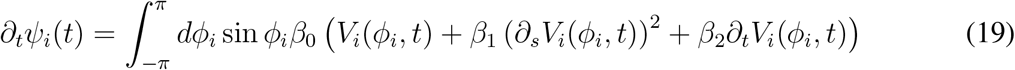

As the system remains invariant when flipping *ϕ* → −*ϕ*, the square of the spatial derivative should be considered. Also, due to the binary nature of the visual field, higher order, and cross, derivatives do not contain more information. Similarly, rising the parameters of the visual field to a higher exponent yields the same function. Finally, we have proposed to neglect the temporal derivative to observe only effects that are instantaneous in time.

### 2 Stable position

In order to understand where the two fish could be in relation to each other, it is important to define the transition between the zone of interaction, the stable state of the system. The relative position of two individuals, *f*_*i*_ and *f*_*j*_, respective to each other, is conserved if they are moving in the same direction *ϕ*_*i*_ = *ϕ*_*j*_, with the same speed *v*_*i*_ = *v*_*j*_. It comes trivially that *∂*_*t*_*v*_*i*_ = 0 if *ϕ* = *kπ* + *π/*2 with *k* integer and that *∂*_*t*_*ψ*_*i*_ = 0 if *ϕ* = *kπ*. Other solutions are then defined by

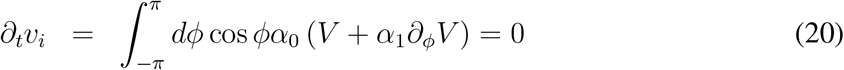

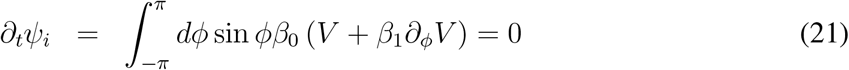

For a pairwise interaction this become

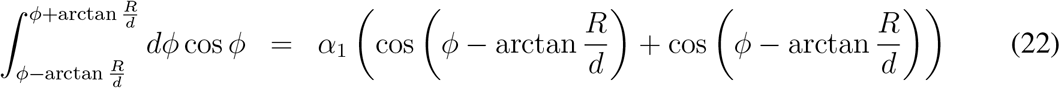

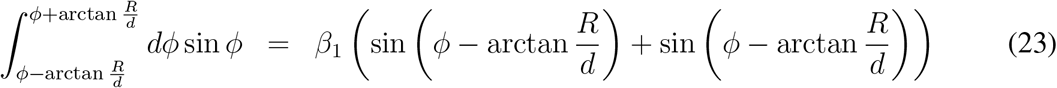

and

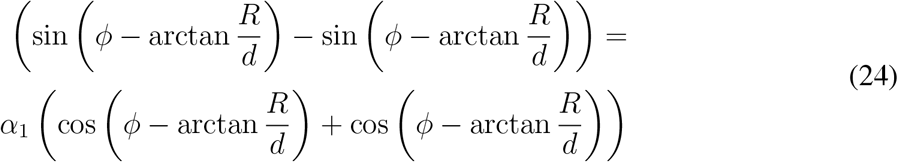

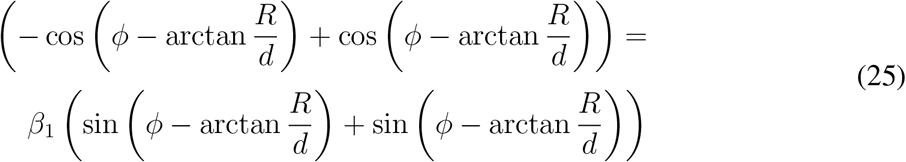

which can be simplified as

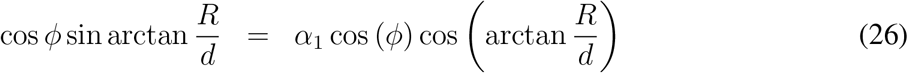

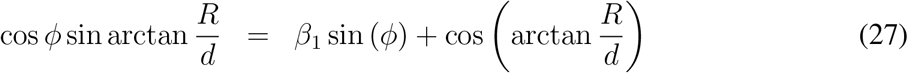

and finally

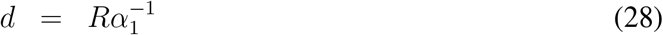

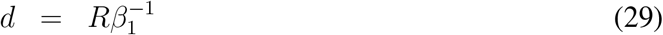

The solution defined a circle of radius 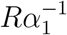 around the individual (Figure 5.A and B). In order to get repulsion for more than one body length 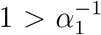 and 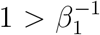. If both lengths are similar a stable position is defined on a ring around the individual (Figure 5.C). The solution being on a circle, the position is stable for both individual and a stable polarization state should be possible. If both lengths are different, only two stable positions are then found, *ϕ* = *nπ/*2. A line where each individual is moving perpendicularly to the main axis of the swarm has been easily observed in the simulation.

**Figure 5:**
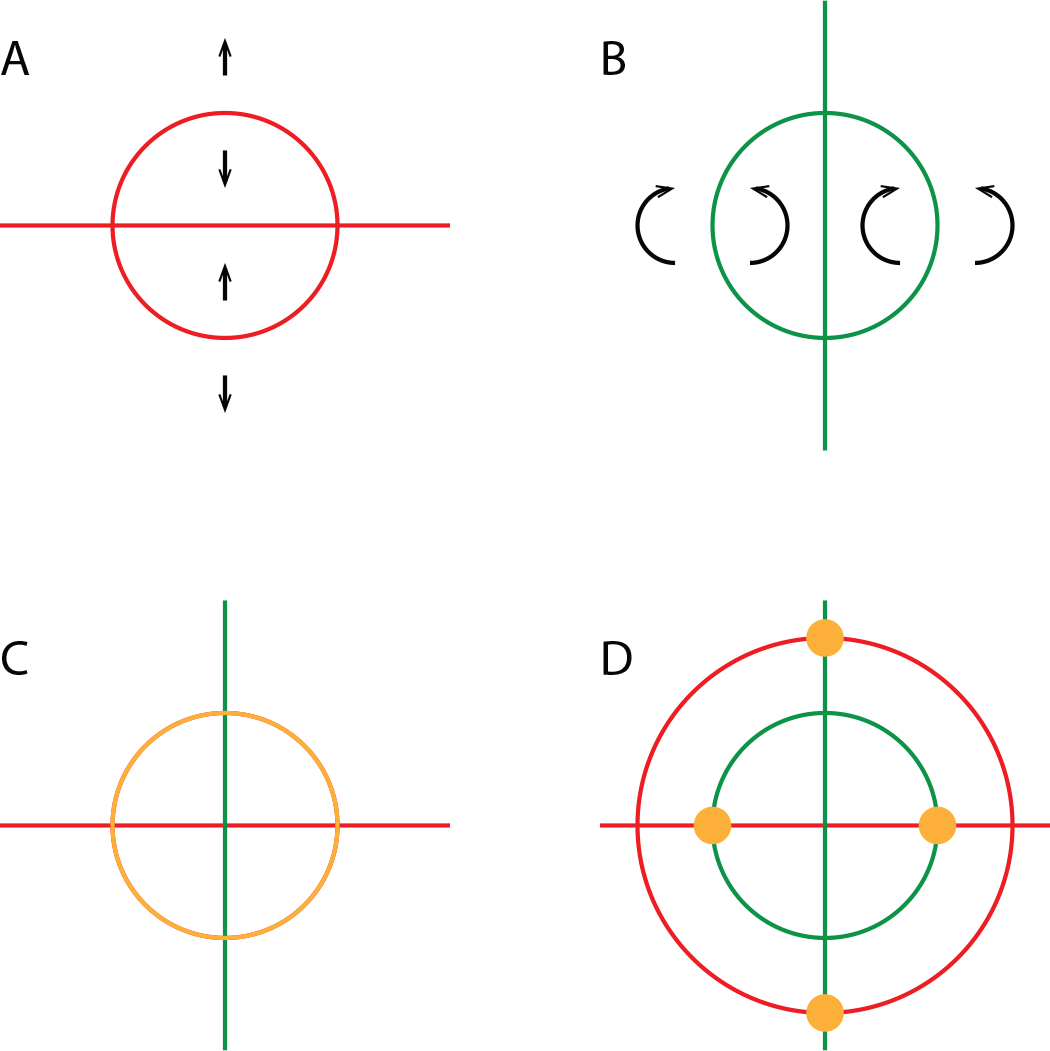
Stable Solution defined for the speeding force in red (A) and the turning force in green (B). Each stable solution is defined by the combination of a line and a circle. The solutions inside the circle are unstable, and only the solutions on the circle are stable. C. If the distance of the stable circle is the same for the turning force and the speeding force a ring of stable solution is defined (in orange). If both distance are different, there exists 5 stable positions (in orange + the center of the circle) but only the two solutions the further from the center are stable.

### 3 Numerical Simulations

A series of numerical simulations has been realized for *N* = 2, 3, 4, 5, 6, 7, 8, 9, 10, 20, 50, 100, 200, 500. Only a subset of this simulation is detailed here (*N* = 2, 10, 20, 50, 100), but video of the movements can be seen at^17^. Each individual agents is a disk of radius *R* = 0.5, self-propelled with a velocity *v* = 1, and the speed relaxation rate is *γ* = 0.1. The time step between two updates of the velocity is ∆*t* = 0.1 while the whole simulation runs until a time *T*_*end*_ = 1000. In all simulations *α*_1_ = *β*_1_ so that the equilibrium distance in the front-back and left right direction are the same. Two different values are used so that *α*_1_ = *β*_1_ = 0.1 and 0.02 accounting for equilibrium distance 5 and 25. For each set of parameters, range of values of *α*_0_ and *β*_0_ have been explored to study the influence of different sensitivity to acceleration and turning rate spanning multiple order of magnitude, *α*_0_ = 0, 0.01, 0.02, 0.05, 0.1, 0.2, 0.5, 1, 2, 5, 10 and *β*_0_ = 0.01, 0.02, 0.05, 0.1, 0.2, 0.5, 1, 2, 5, 10.

Due to the geometry of the system considered here, the individual agents, it was possible to avoid techniques such as ray-casting. Instead the visual field of each individual, *V*_*i*_, is considered as a linear function of *ϕ*, from *−π* to *π*, constituted of 16384 elements, so that ∆*ϕ* = *π/*8192. *V*_*i*_(*ϕ*) is initialized at 0 at each time step. The visual field is then updated by considering the relative position of each individual *j*, in the referential of the focal individual *i* oriented in the direction of the movement 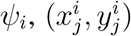. The position in the visual field is recovered by computing the relative angle of each individual 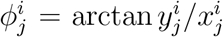, with the extension 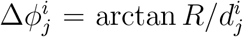, where 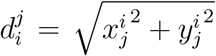. The values of the visual field are the updated so that *V*_*i*_ (*ϕ*) = 1 in the range 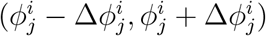. As the visual field is binary, there is no need to considered supplementary calculation to account for occlusion. The velocity and turning rate of each individual at time *t*+∆*t* is then updated accordingly

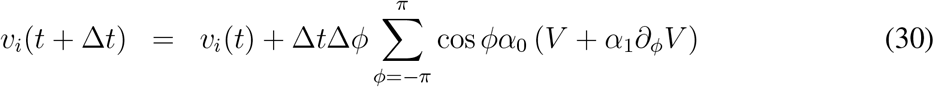

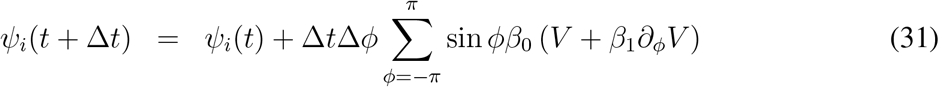

The minimal distance observed, *d*_*m*_*in* is measured by taking the minimal inter-individual distance at each time step and updating for all the time of the simulation

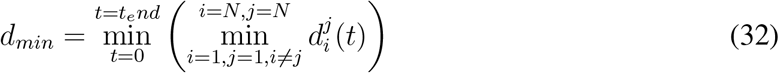

If *d*_*m*_*in* < 1, then collisions are observed and the computation is stopped.

The mean polarization, *p* is measured as the average polarization between all individuals at each time step, averaged on the last 10% of each simulation, so in the temporal range *t* = (900, 1000),

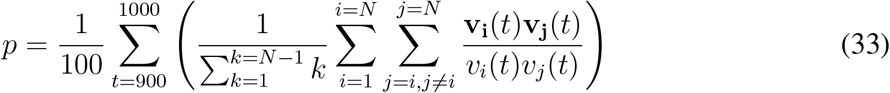

Finally the average closest neighbor distance is measured, *d*_*mean*_ as the average 10% of each simulation, so in the temporal range *t* = (900, 1000), of the minimal inter-individual distance observed at each time step

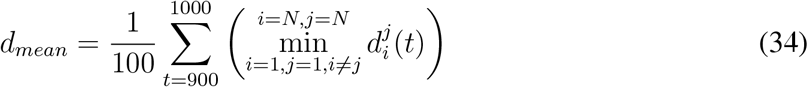

## Aknowledgements

RB wants to thank all the people who gave support during various part of the research process, starting with the origin of the project as a multiplayer virtual reality videogame, this includes Iain Couzin, Olivier Dauchot, Guy Theraulaz and Hunter King. RB acknowledges funding by the Young Scholar Fund of the University of Konstanz. PR acknowledges funding by the Deutsche Forschungsgemeinschaft (DFG, German Research Foundation) under Germany’s Excellence Strategy – EXC 2002/1 “Science of Intelligence” – project number 390523135, as well as through the Emmy Noether program, project number RO4766/2-1.

